# Non-destructive X-ray tomography of brain tissue ultrastructure

**DOI:** 10.1101/2023.11.16.567403

**Authors:** Carles Bosch, Tomas Aidukas, Mirko Holler, Alexandra Pacureanu, Elisabeth Müller, Christopher J. Peddie, Yuxin Zhang, Phil Cook, Lucy Collinson, Oliver Bunk, Andreas Menzel, Manuel Guizar-Sicairos, Gabriel Aeppli, Ana Diaz, Adrian A. Wanner, Andreas T. Schaefer

## Abstract

Maps of dense subcellular features in biological tissue are the key to understanding the structural basis of organ function. Electron microscopy provides the necessary resolution, yet - as electrons penetrate samples for only a few 100s of nm - requires physical sectioning or ablation, which strongly challenges anatomical investigations of entire organs such as mammalian brains. As demonstrated for the engineering and physical sciences, X-ray nanotomography represents a promising alternative for ultrastructural 3d imaging without physical sectioning^1–15^. Leveraging the high brilliance of 4th generation synchrotron X-ray sources, it has the potential to non-destructively image mm³-sized samples at ultrastructural resolution within a few days^16^. A fundamental barrier to application to the life sciences is that, when irradiated with high-intensity X-rays, biological samples deform and ultimately disintegrate, prohibiting reaching sufficient resolution. Here, we introduce a combination of engineering solutions which defeat this barrier for X-ray ptychography^17^, a coherent diffractive X-ray imaging technique. The solutions include a cryogenic sample stage with high stability, high-precision interferometric positioners and tailored non-rigid tomographic reconstruction algorithms^18^. Furthermore, adapting an epoxy resin developed for the nuclear and aerospace industry, we demonstrate radiation resistance to X-ray doses exceeding 10^10^ Gy. The resulting sub-40 nm isotropic resolution makes it possible to densely resolve axon bundles, boutons, dendrites and reliably identify synapses without physical sectioning. Moreover, we validated the X-ray technique using the current gold standard, namely focused ion beam scanning electron microscopy (FIB-SEM)^19,20^ to demonstrate intact ultrastructure in tissue volumes first imaged by X-rays. This unlocks the potential of X-ray tomography for high-resolution tissue imaging, coinciding with the transformative advancements of next-generation synchrotrons worldwide^21^.

## Introduction

Current gold-standard methods for dense tissue reconstruction rely on volume electron microscopy^22–29^. As electron penetration depth into tissue is limited to at most a few 100 nm^1,30^, obtaining 3-dimensional information requires tissue sectioning or milling either before preparing samples for imaging^25–27^ or during image acquisition^31–33^. Such workflows present significant challenges for reliable continuous image acquisition, with the risk of information loss during month-long experiments, and experimental analysis which requires reconstruction and alignment of multimodal datasets.

X-rays can penetrate samples of several mm to cm, allowing for largely non-destructive imaging of tissues^1,30^. Laboratory X-ray sources can resolve cell bodies and even large neurites^3^ but their limited brilliance makes high-resolution imaging extremely time-consuming. Large scale synchrotron facilities can provide high-brilliance X-rays at wavelengths in the Ångstrom range, and are therefore driving the development of imaging methods capable of providing 3D resolution down to 10s of nanometres for hard, inorganic nanostructures with high density contrast^34–38^. Application of these methods to biological imaging has enabled reliable identification of various biological features including blood vessels, cell bodies and large neurites^4–15^. More recently, thin neurites were identified at a resolution of ∼100 nm using nano-holotomography^2,39^. Fundamentally, however, the resolution in X-ray tomography scales with the 4th power of the radiation dose, i.e. to increase the resolution by a factor of 5 in the same sample, the X-ray dose needs to be increased by a factor of 5^4^=625-fold. X-ray imaging of soft biological tissue was previously thought to be limited by tissue contrast and by the radiation dose that can be employed before the tissue is damaged^1,30,40^, and it remained unclear whether key ultrastructural features could be examined with X-rays at sufficient resolution.

To overcome the limited X-ray image contrast inherent to soft tissues, we combine heavy-metal staining protocols developed for volume electron microscopy^41^ with high-resolution ptychographic tomography^34,42^. Radiation damage is minimised by imaging samples under cryogenic conditions using a recently developed cryogenic ptychographic tomography instrument^10,43^ and by embedding them in a newly identified highly radiation-resistant resin^44–46^. Residual sample deformations will introduce inconsistencies between the acquired projections, for which we compensate computationally, in order to avoid loss of 3D reconstruction quality^18^. By combining these techniques, we obtain an isotropic 3D resolution of 38 nm for neural tissue, sufficient for resolving synapses, while simultaneously maintaining tissue integrity.

## Results

X-ray ptychography is performed by scanning a sample across a confined, coherent beam a few µm in diameter and measuring diffraction patterns in the far field (**Fig. 1(a)**)^17^. Using iterative reconstruction algorithms^47,48^, these diffraction patterns (**Fig. 1(a)**) are then used to produce phase images, which are proportional to the projected electron density of the sample (**Fig. 1(b)**). To perform ptychographic X-ray computed tomography (PXCT), the projections are acquired at multiple sample rotation angles from 0 to 180 degrees, followed by tomographic reconstruction of a 3D sample volume (**Fig. 1(c)**). Resolution in X-ray ptychography is limited not by the beam size but by the angular acceptance of the detector, which determines the maximum observable spatial frequencies, and the applied X-ray dose, which determines the signal-to-noise ratio of the spatial frequencies within the recorded diffraction patterns. Since the imaging geometry can be adjusted relatively easily, radiation damage and weak intrinsic scattering of biological tissues are the main factors limiting the attainable resolution^16,30,40^. We therefore employed a recently developed cryo-tomography stage^43^ to cool samples to <95 K and to reduce the structural changes caused by ionising radiation, in combination with a staining and embedding protocol to maximise contrast. Additionally, we applied a new non-rigid tomography reconstruction algorithm, designed to compensate for radiation-induced deformation of the sample^18^. In the following, we present tomograms from mouse brain tissue (the radiatum layer of hippocampus, the external plexiform and glomerular layers from the olfactory bulb, and upper layers of neocortex as structures rich in neurites and dendritic shaft and spine synapses) at different radiation doses.

**Figure 1:**
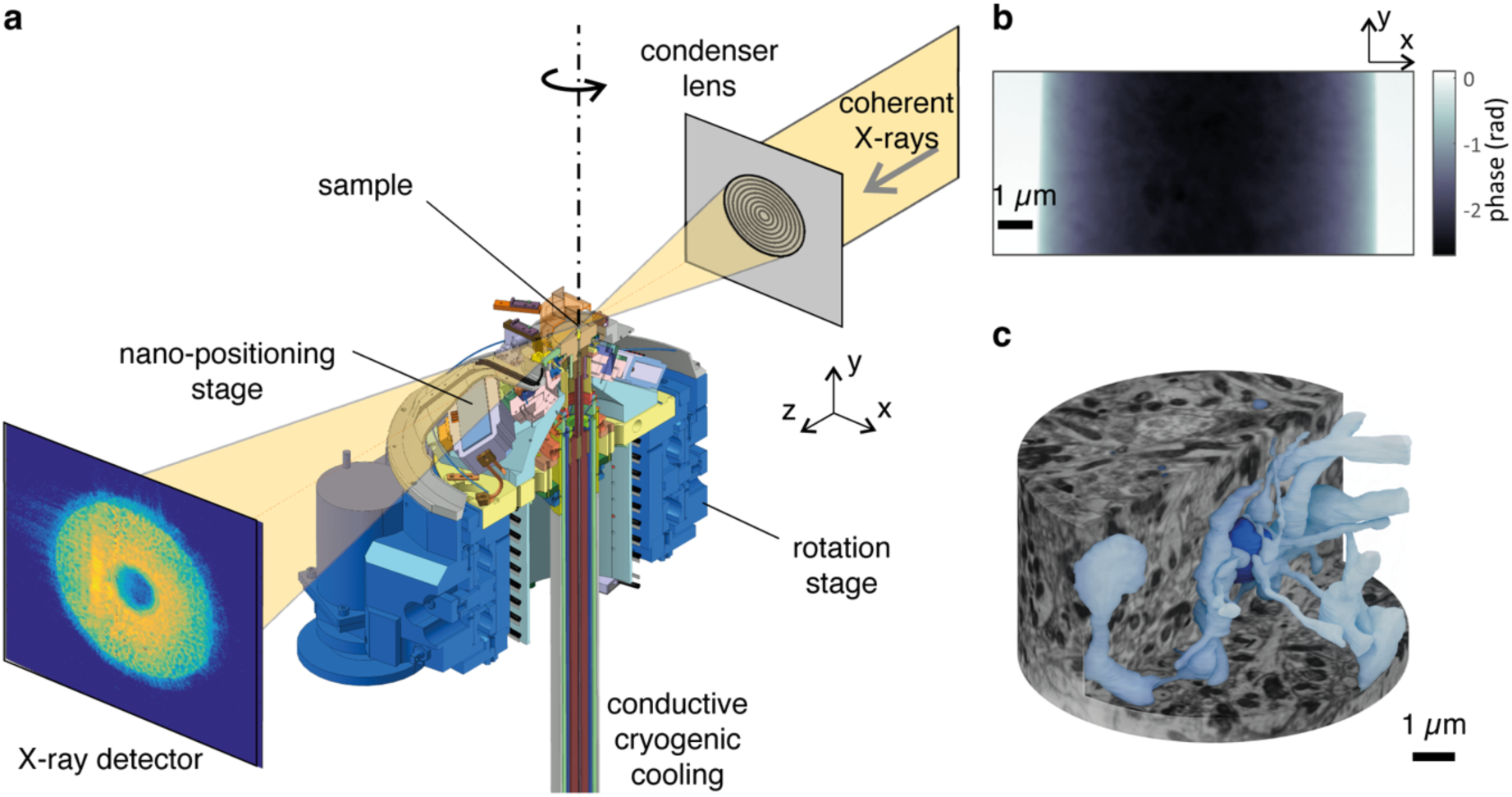
X-ray ptychography of brain tissue. **(a)** Ptychography setup; top right: incoming coherent beam is focused by a lens to define the illumination on the sample; center: cryogenic stage for scanning with 10nm accuracy with rotation capability; bottom left: example diffraction pattern recorded by the X-ray detector. **(b)** Single reconstructed phase projection of a Resin E (TGPAP)-embedded mouse brain tissue sample extracted from the external plexiform layer in the olfactory bulb, obtained using a monochromatic 6.2keV beam under cryogenic conditions at cSAXS. **(C)** 3D rendering of the reconstructed tomogram containing the projection shown in (b) (1190 projections, total absorbed dose 2.5e9 Gy). A part of the volume has been cut to show the segmentation of a neurite (dark blue) and some of its presynaptic partners (light blue), based on the X-ray tomogram dataset.

To assess what brain structures PXCT can resolve, we stained external plexiform layer tissue from mouse olfactory bulb with a ROTO stain^41^ embedded it in a ‘hard Epon’ resin^49^ (hard epon resins A and B, **Supp Fig. 5.1, Supp Table 3**, see resin descriptions in the section Materials & Methods) and prepared pillars with a diameter of 10-30 µm. At a dose of 1.8×10^7^ Gy we were able to readily resolve dendrites (**Fig. 2(a)**, **Supp Fig. 2.2(a1)**); using Fourier shell correlation (FSC, solid line in **Supp Fig. 2.2(c)**)^50,51^ we estimated an isotropic 3D resolution of 84 nm (**Supp Fig. 2.2(c1, d)**). Cryo-cooling is thought to reduce structural changes that limit resolution in X-ray imaging due to radiation damage^30,52^. Consistent with this notion, acquired tomograms did not show any indication of significant blurring or other signs of structural changes due to radiation damage. Irradiating a sample with an increased radiation dose of 8.4×10^7^ Gy resulted in a 3D resolution of 69 nm (**Supp Fig. 2.2(a2, c2, d)**). However, contrary to theoretical predictions^16^, an even higher dose of 3.8×10^8^ Gy did not enable an enhanced resolution (70 nm; **Supp Fig. 2.2(a3, c3, d)**). In particular, the sample deformed, presumably due to “softening” of the embedding resin, as seen when comparing low resolution tomograms acquired with ∼2.6×10^8^ Gy intermittent irradiation (**Supp Fig. 2.1**). As a result, the resolution of the combined tomogram decreased based on visual blurring in the 3D volume and the decrease of the correlation at several spatial frequencies, as determined by FSC analysis **Supp Fig. 2.2(a3, c3)**. To take these sample changes into account, we applied a non-rigid tomography reconstruction approach that estimates the deformations between the subtomograms and includes them into the tomographic reconstruction ^18^(**Supp Fig. 2.1**). This deformation correction method requires tailored data acquisition where the whole angular range from 0 to 180 degrees is split into multiple equally spaced angular ranges to acquire multiple subtomograms. In doing so, each subtomogram provides a low-resolution snapshot of the sample at a given time point. By calculating the 3D optical flow between these subtomograms, the deformation fields can be estimated across a range of time points and interpolated to describe the changes between all collected projections. Accounting for radiation-induced changes in this manner recovered higher spatial frequencies (as seen in **Supp Fig. 2.2(b)** and from the FSC in **Supp Fig. 2.2(c)**) which translated into an improved 3D resolution of 49.7 nm (**Fig. 2(b,d)**, **Supp Fig. 2.2(b3,c3,d)**), i.e., close to the theoretical predictions of resolution improving with the fourth power of dose^40^. The use of non-rigid tomography consistently improved the expected scaling between the radiation dose and attainable resolution across the measured samples (**Fig. 2(d), Supp Fig. 2.2(d)**). Measured at a temperature of ∼90 K, tissue was radiation-stable at doses up to 6×10^8^ Gy (**Supp Fig. 2.4**) when embedded with either of the traditional EM-embedding resins EMbed812 or Epon812^53^ (see methods).

**Figure 2:**
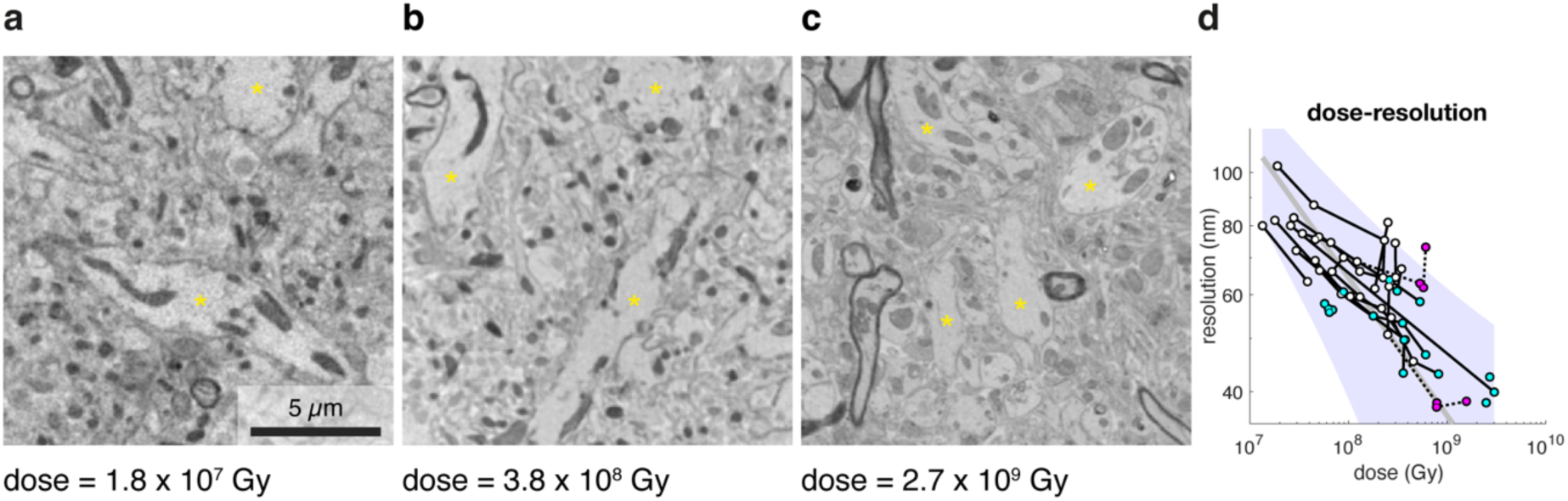
Dose-dependence of resolution for X-ray ptychography of brain tissue. **(a-c)** Example 2D slices from 3D non-rigid reconstructions of tomograms acquired at increasing radiation doses of samples embedded in resin A (EMbed812) (a-b) and resin E (TGPAP) (c). Yellow asterisks indicate dendrites of projection neurons in all three images. **(d)** Fourier shell correlation (FSC) resolution for all tomogram series of all samples reported, imaged at different doses. Tomograms (circles) obtained of the same sample at multiple doses form a series and are linked with a black edge. The colour of the tomogram nodes indicates whether the tomogram had no radiation-induced damages (white), had contained, globular and compact (<100 nm) low-absorbing regions compatible with reconstruction and analysis (cyan) or a diffuse mass loss (magenta). See **Fig. 5** for a detailed description of the different types of observed radiation damage. A power function *FSC_resolution = a/dose^(¼)* (grey line) fitted to all white and cyan data points along its 95% prediction intervals with observation, non-simultaneous bounds (purple shaded area) is shown.

Using this improved approach with imaging ROTO-stained samples at cryogenic temperatures with non-rigid tomographic reconstructions we obtained 3D tomograms for ROTO-stained and either Epon812 or EMbed812-embedded tissue from a variety of brain regions, including olfactory bulb external plexiform layer, glomerular layer, hippocampus stratum radiatum, and layer 2-3 of somatosensory cortex (**Fig. 3**). This allowed us to visualise e.g. neurites, mitochondria, myelin, or nuclei (**Fig. 3**).

**Figure 3:**
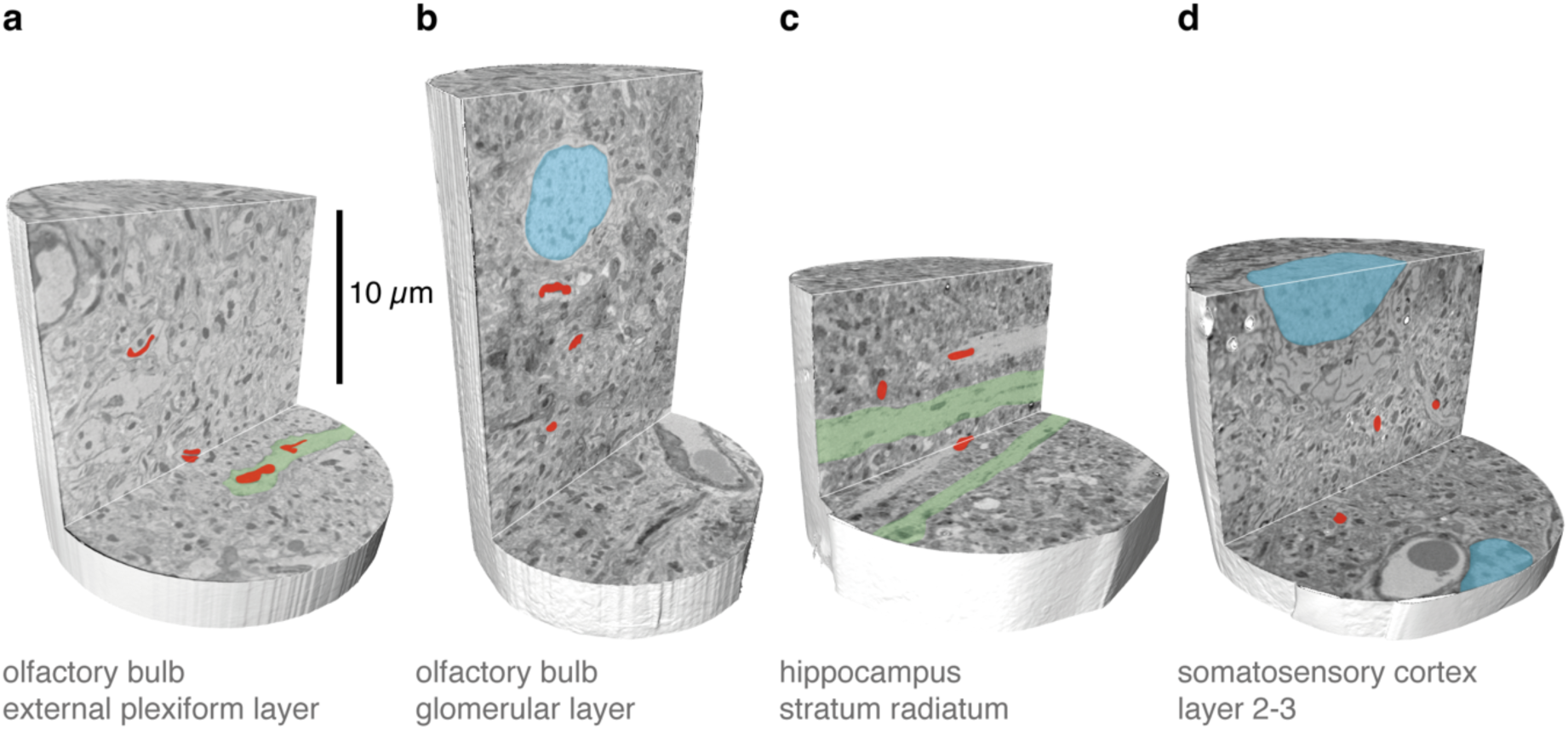
Subcellular features detectable with X-ray ptychography in different brain tissues. **(a-d)** Cutout views from X-ray ptychography tomographic reconstructions for different tissues (imaged using radiation doses of 3.8 x 10^8^ Gy, 3.8 x 10^8^ Gy, 2.2 x 10^8^ Gy, 1.9 x 10^8^ Gy). The respective tissue samples were embedded in resin A (EMbed812) (a-c) and resin B (Epon812) (d). Subcellular features including dendrites (green), nuclei (blue) and mitochondria (red) are resolved.

A critical feature for connectomics is the detection of synaptic contacts (**Fig. 4(c)**). Thus, we assessed whether PXCT would allow the identification of such structures. Indeed, for a dose of 3.8×10^8^ Gy in Epon-embedded olfactory bulb samples, structures resembling synapses became apparent (**Fig. 4(a)**). PXCT being a largely non-destructive technique, we subsequently processed the same specimen using focused ion beam milling combined with scanning electron microscopy (FIB-SEM). This allowed us to obtain insights into potential ultrastructural alterations induced by the X-ray irradiation as well as obtain a “ground truth” for structure identification and further quantification. The FIB-SEM reconstruction showed a mild curtaining effect that might be the consequence of radiation-induced changes to the embedding medium (**Supp Fig. 4.1**). However, the overall ultrastructure was not noticeably impacted by prior X-ray radiation dosage of ∼4×10^8^ Gy (**Fig. 4(b)**). In particular, membranes were left intact, as were subcellular features such as mitochondria, endoplasmic reticula or vesicles (**Fig. 4(b)** and **Supp Fig. 4.1**). Aligning the FIB-SEM with the PXCT dataset allowed us to directly compare structures identified in PXCT with the ground truth data obtained from FIB-SEM (**Fig. 4(a,b,c)**). To quantify the reliability of synapse identification, we performed two approaches: Firstly, we traced individual dendrites in both datasets (**Fig. 4(d))** and performed independent manual labelling of synapses in both datasets (**Fig. 4(d)**). More than two-thirds of synapses identified in the FIB-SEM dataset were recognized in a blind analysis of PXCT images (225/338 synapses, 70±6%, n=5 dendrites **Fig. 4(d,e)**). Additionally, to obtain true estimates of precision and reliability, three annotators were presented with randomly sampled 1 µm^3^ volumes of the FIB-SEM or PXCT datasets and asked to indicate whether or not a synapse was present in the volume with a confidence score (**Supp Fig. 4.2**). Using the FIB-SEM dataset as ground truth again, PXCT annotation showed a recall of synapse presence of 60% with a precision of 80% (**Fig. 4(f)**, **Supp Fig. 4.2**).

**Figure 4:**
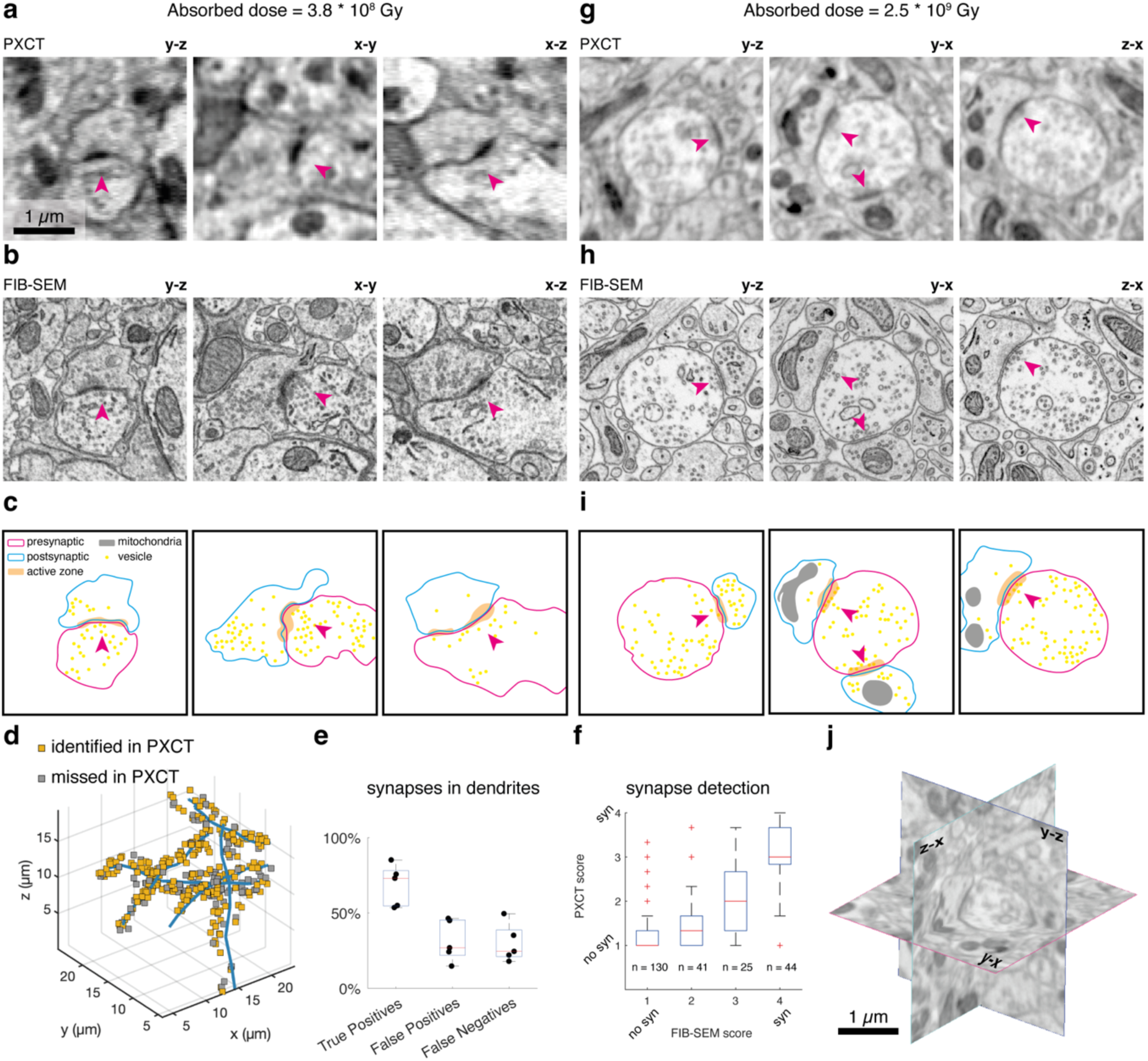
Quantitative evaluation of synapse detectability with correlative X-ray ptychography and FIB-SEM. **(a)** Example PXCT images of a dendrodendritic synapse (magenta arrowheads) from the external plexiform layer in 3 orthogonal reslices, acquired from a tissue sample embedded in resin A (EMbed812). Note the prominent synapse in the center. **(b)** Same region in (a) from the corresponding FIB-SEM dataset. **(c)** Segmented features of the neurites establishing the synapses shown in (b). **(d-e)** 3 dendrite branches traced in PXCT and FIB-SEM with synapses highlighted that were detected in both (orange squares, n=225) or in FIB-SEM only (grey squares, n=113). **(f)** In an independent analysis, presence or absence of synapses was evaluated in n=240 randomly selected 1 µm^3^ subvolumes in both FIB-SEM and PXCT volumes. The box plot indicates the average score given to the distinct ROIs, binned by their EM scores. **(g)** Example PXCT images of a dendrodendritic synapse (magenta arrowheads) from the external plexiform layer in 3 orthogonal reslices, acquired from a tissue sample embedded in resin E (TGPAP). **(h)** Same region in (f) from the corresponding FIB-SEM dataset. **(i)** Segmented features of the neurites establishing the synapses shown in (h). **(j)** 3-dimensional view of the PXCT scene shown in (g).

Increasing resolution further would benefit in particular automated reconstruction methods ^54,55^ but this would require an even higher X-ray dose. Epon-embedded samples started displaying deformations consistent with mass loss possibly compromising ultrastructural features. In agreement with previous work^16,40^, significant radiation damage such as widespread mass loss was consistently observed at doses > 6×10^8^ Gy (**Fig. 5(b,c,d)**, **Supp Fig. 2.4**). In order to improve radiation resistance, we turned to resins previously developed for nuclear reactors or the aerospace industry^44–46^, optimised for stability in the presence of ionising radiation. Screening a battery of such resins, we identified a low viscosity epoxy with good infiltration properties consisting of triglycidyl para-aminophenol (TGPAP) (**Fig. 5(e)**), a triglycidyl ether of para-aminophenol. Like Epon812, TGPAP monomers are tri-functional, i.e. they contain three epoxy groups, but additionally also a benzene ring and a tertiary amine group. For curing, we used the 4,4’-diaminodiphenyl methane (DDM), a well-known hardener with two primary amine groups that react with the epoxy groups of TGPAP (**Supp Fig. 5.1, Supp Tables 2, 3**). The three epoxy groups per molecule allow for increased crosslinking during curing, leading to superior mechanical strength, chemical and radiation resistance, as well as thermal stability. Moreover, the benzene rings in both the monomer and the hardener are thought to further increase radiation resilience. To assess radiation resistance, we exposed samples embedded in Epon812 or TGPAP to a broad bandwidth beam with an average energy of 25 keV, generated by a bending magnet at ESRF’s beamline BM05^56^. While Epon samples disintegrated at <10^9^ Gy (based on measurements with a monochromatic 6.2 keV beam at SLS), TGPAP samples showed little sign of mass loss or compromise of cellular features even with prolonged exposure (**Fig. 5(f)**, **Supp Video 5.1**). When we reconstructed PXCT tomograms from Epon812 at different total doses we noticed visible damage and mass-loss in the irradiated areas once total dose exceeded ∼6×10^8^ Gy (**Fig. 5(c,d)**). Using TGPAP for embedding (with the same ROTO staining and mouse olfactory bulb tissue) we found no indication of mass loss and only localised sub-micron punctae that seemed to emerge with a total dose of >10^9^ Gy but did not impact on relevant tissue features (**Fig. 5(g,h)**, **Supp Fig. 5.2**). Importantly, TGPAP tissue penetration is comparable to commonly used Epon and the ultrastructure is well preserved (even after irradiation with a total dose exceeding 2×10^9^ Gy) as verified by FIB-SEM analysis (**Fig. 5(i,j)**).

**Figure 5:**
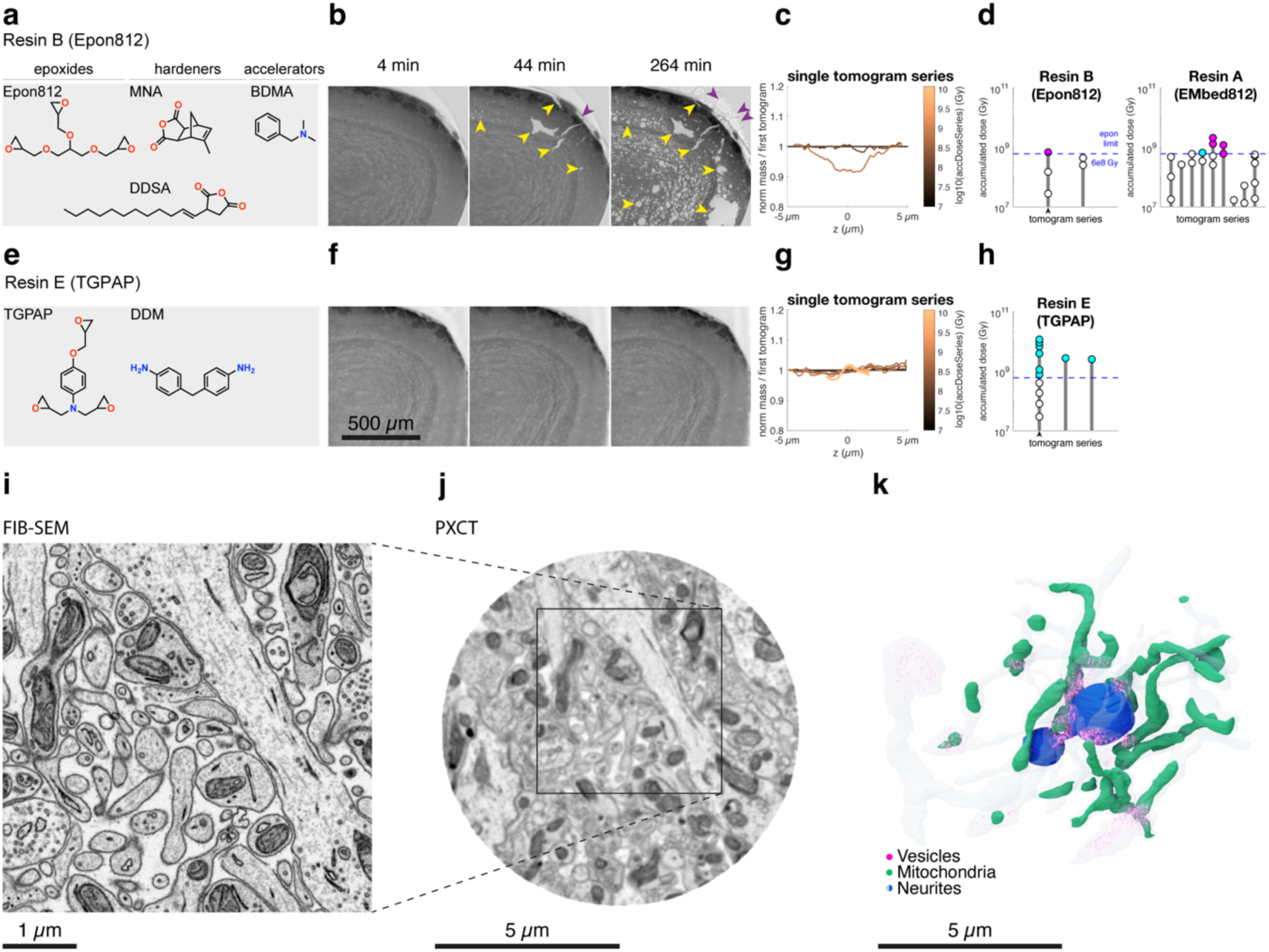
A ‘tough’ epoxy resin for increased radiation resistance. **(a)** Chemical structure of the monomers used to polymerize the two most representative resins explored in this study, resin B (Epon812-MNA-DDSA-BDMA, later referenced as ‘Epon812’ for simplicity). **(b)** Radiation exposure tests on stained olfactory bulb samples embedded in resin B (Epon812), performed at room temperature and ambient pressure using a filtered polychromatic X-ray beam with average energy of 25 keV (ESRF BM05). Assessment of radiation effects is based on phase contrast microtomographic imaging. Note the widespread resin degradation within (yellow arrowheads) and outside (purple arrowheads) the embedded tissue. **(c-d)** Mass variation across the vertical direction in a series of PXCT tomograms obtained from the same sample embedded in resin B (Epon812) reveals a decrease in mass in the central region that was more exposed to radiation in the latter tomograms. This effect was observed in almost all tomogram series after the accumulated dose exceeded 6.8e8 Gy, both in samples embedded in resin B (Epon812) (d, left) as well as in those embedded in resin A (EMbed812) (d, right). **(d)** Stacked doses of all tomograms of each series, with tomograms affected by this artefact coloured in magenta. Tomogram series reported in (c) is indicated by a black arrowhead. **(e-h)** Same as (a-d), for samples embedded in the radiation ‘tough’ Resin E (TGPAP-DDM, later referenced as ‘TGPAP’ for simplicity). The disaggregated mass loss effect described in (b) was never observed in samples embedded with TGPAP (f), which instead showed mass loss in very compact and isolated <100 nm wide foci, not affecting the integrity of the specimen (g-j) and allowing for faithful reconstruction of the tomograms even after absorbing beyond 1.2e10 Gy. This effect was observed in all TGPAP tomogram series (h, affected tomograms coloured in cyan). **(i-j)** The infiltration quality and the ultrastructural integrity for the TGPAP embedded samples was assessed at high resolution using FIB-SEM (i) after PXCT (j). **(k)** Some neurites segmented using the PXCT data only. A dark blue neurite receiving synaptic contacts from transparent pale blue neurites is shown alongside mitochondria (green) and vesicle clusters (magenta).

Embedding heavy-metal stained tissue in TGPAP therefore allowed us to significantly increase the cumulative radiation dose for PXCT. This in turn enabled us to further increase the resolution to 38 nm (**Fig. 5(j)**, **Fig. 2(c,d)**, **Supp Video 5.2**). The resulting resolution was sufficient to resolve and segment vesicle clusters and synaptic densities (**Fig. 5(k)**, **Supp Video 5.3**). Importantly, FIB-SEM milling confirmed that the ultrastructure was intact after irradiation with a dose exceeding 10^10^ Gy. Together, this indicates that using newly identified radiation-resistant embedding materials with heavy metal staining and non-rigid tomographic reconstruction methods, X-ray nanotomography can provide sufficient 3D resolution to identify key connectomic features.

## Discussion

Here, we have demonstrated the potential for synchrotron X-ray nanotomography to image at resolutions sufficient to reliably detect key ultrastructural features such as synaptic contacts. While employing sub-nm wavelengths, resolution in X-ray imaging of biological soft tissue is generally limited by contrast. Furthermore, the softness of embedding materials raises susceptibility to radiation damage, which ultimately limits the maximum attainable resolution. When heavy metal stains are employed, they increase contrast and X-ray absorption. Absorption, however, underlies radiation damage^1,30,40^. At the Swiss Light Source, we employed the tomography nano cryo stage (OMNY), an instrument that uses an interferometer-guided cryogenic stage optimised for PXCT measurements^43^. Moreover, we identified an epoxy resin with superior radiation resistance^44–46^. Together this allowed us to mitigate the effects of radiation damage and significantly improve the resolution of X-ray imaging of biological tissue to the extent that key structures such as vesicles or synaptic densities can be densely detected in 3D.

At a dose of 4×10^8^ Gy we could detect radiation-induced changes, e.g. sample expansion, and some changes such as curtaining effects and small deformations were visible outside the imaged area in the downstream FIB-SEM analysis (**Supp Fig. 2.2**, **4.1**). However, none of these alterations resulted in any significant ultrastructural changes. We have also shown that non-rigid tomography reconstruction algorithms^18^ can largely compensate for radiation-induced deformations and improve resolution by modelling the tissue expansion explicitly (**Fig. 2**, **Supp Fig. 2.1**). Moreover, the OMNY stage, which we have developed previously^43^, allows for cooling to liquid helium temperatures. Cooling to such even lower temperatures might further reduce radiation damage, allowing for additional increases in radiation dose and resolution.

Embedding and staining protocols have played a key role in the development and improvement of volume EM techniques^41,57–62^. Epon embedding provides excellent mechanical properties for thin serial sectioning^63^ but is prone to radiation damage^64^ (**Fig. 5(b,c,d)**, **Supp Fig. 5.2**). To enhance radiation resilience, we therefore turned to the aerospace and nuclear industry that critically depends on epoxies resistant to ionising radiation. We identified one resin (TGPAP-DDM, referred to as ‘TGPAP’ and ‘Resin E’ for simplicity) that penetrated tissue well, preserving ultrastructure, and showed significantly improved radiation resistance. In fact, we could not observe substantial mass loss or significant damage to tissue samples at radiation levels that caused macroscopic holes and widespread damage in Epon812 embedded samples (**Fig. 5(f)**, **Supp Video 5.1**). Therefore, we call the TGPAP polymer ‘tough resin’. Theoretical estimates for flux and dose constraints for X-ray tomographic imaging indicate that such dose resistance will suffice for whole-mouse brain X-ray tomographic imaging at sub 20 nm resolution^16^.

While not resulting in tissue disintegration or ultrastructural damage, irradiation results in localised tissue alterations as well as global deformations, likely due to thermal expansion or limited impact on chemical bonds and e.g. local gas formation (**Supp Fig. 5.2**). Ultrastructure (e.g. membranes, protein clusters) seem to be largely unimpacted (**Figs. 4(b,h)** and **5(i)**); however, such deformations result in blurred tomographic reconstructions, limiting resolution (**Fig. 2**, **Supp Fig. 2.2**). We therefore employed non-rigid reconstruction methods that partially mitigate these limitations. Improvements of such algorithms or tomographic acquisition with intermediate, low-dose reference tomograms, might further improve resolution to levels significantly below 20 nm. Correlative volume EM datasets (such as those shown in **Figs. 4(b,h)** and **5(i)**) can provide ground truth for the development of such algorithms. While providing good contrast, heavy-metal stains might in fact not be ideal for X-ray imaging and materials with similar scattering but significantly reduced absorption^65,66^ might further improve sample stability.

Scaling up PXCT to larger volumes presents several challenges. Firstly, in large volumes and at high resolution, X-ray penetration depth is significantly larger than the depth-of-field. With the increased longitudinal coherence enabled by 4th generation synchrotron sources, “multislice” reconstruction methods can, however, explicitly account for such beam propagation effects^67–69^. This not only provides an efficient path to large volume X-ray imaging, but also relaxes the Crowther criterion, reducing the number of rotation angles^70^. Secondly, one practical challenge of using PXCT for neural circuit imaging could be the high photon energy needed to penetrate metal-stained samples of several millimetres, because at these energies it is more difficult to achieve high brilliance at synchrotron sources, and detectors are not as efficient. While more tailored staining protocols might circumvent these limitations, in the near-term, changing the imaging geometry and employing laminography ^35,71–73^ would allow imaging of mm wide slices of e.g. 10-50 µm thickness. Such slices can be produced using hot-knife techniques^74^, currently employed in preparation for FIB-SEM imaging. The versatility of synchrotron X-ray imaging then allows to rapidly acquire overview images followed by repeated targeted high-resolution image acquisition^35^.

Herein, we show that X-ray coherent imaging can densely resolve key connectomic features such as synapses in tissue. Importantly, we demonstrate that biological tissue can be prepared and imaged to withstand radiation dose exceeding 10^10^ Gy, removing the fundamental limitations to employing X-ray tomography for ultrastructural-resolution tissue imaging. The advent of 4th generation, high brilliance X-ray sources will introduce this fundamentally non-destructive, scalable, and highly reliable technique as a widely available, powerful component of the connectomics and wider tissue life sciences toolbox.

## Supplementary Materials

**Supp Fig. 2.1:**
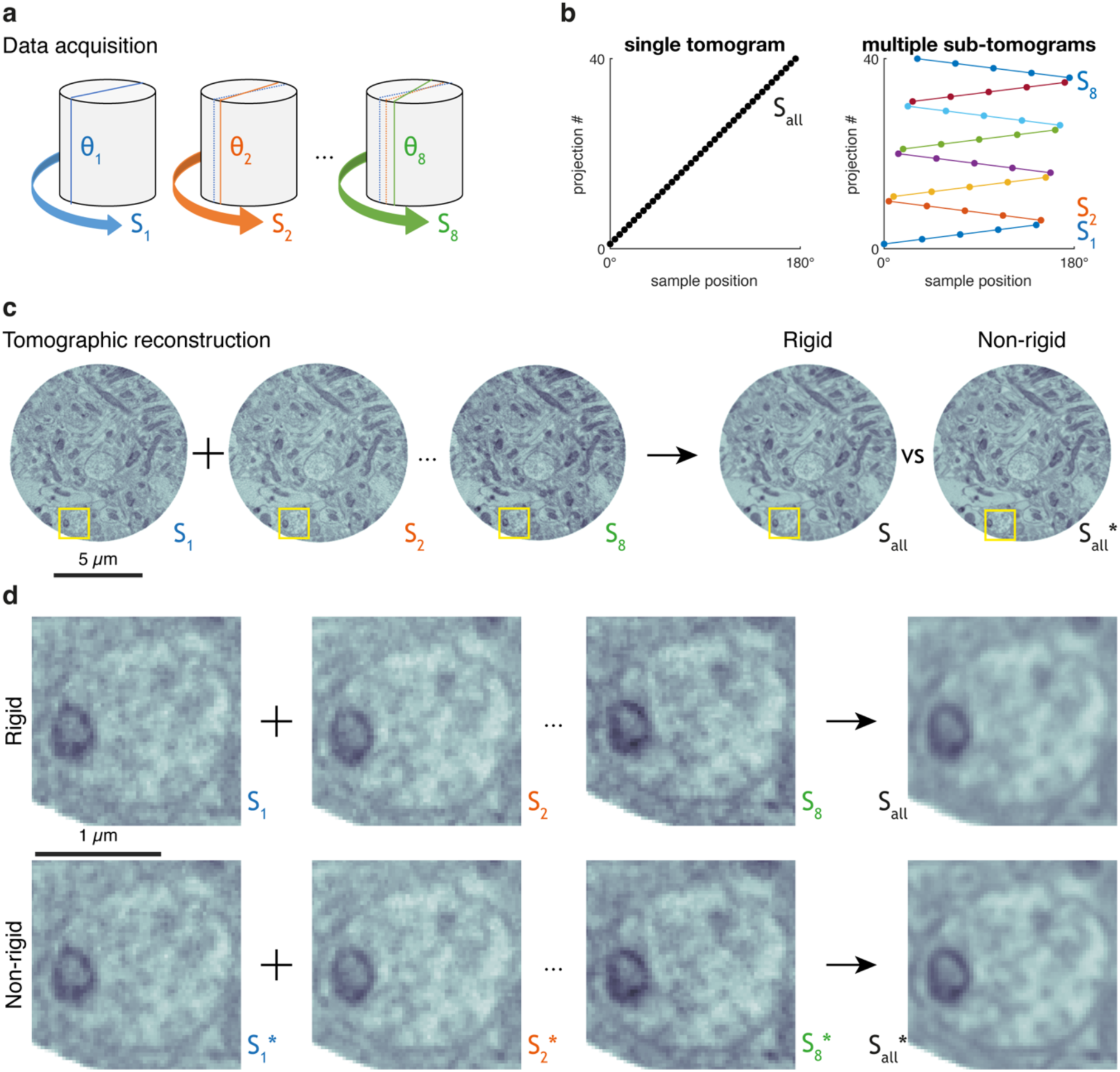
Subtomogram acquisition and non-rigid tomogram reconstruction. **(a-b)** Schematics of a subtomogram acquisition for a mouse brain EPL sample embedded in resin A (EMbed812). Data are acquired in multiple tomograms, each sampling the same angle but starting with an offset from each other (a). This allows for an acquisition algorithm where all desired angles are sampled but in distinct subtomograms (b), each of them enabling independent reconstruction. **(c-d)** Reconstructed subtomograms can later be summed to generate the final reconstruction. This sum can be applied either directly (through a ‘rigid’ approach of all subtomograms S_1_ … S_8_) or after warping all subtomograms to a common space (‘nonrigid’ sum of all warped tomograms S_1_* … S_8_*). Any sample deformations occurring during acquisition will affect a limited subset of subtomograms, and their effect of worsening reconstruction quality is then minimised by the nonrigid reconstruction approach (see insets in d).

**Supp Fig. 2.2:**
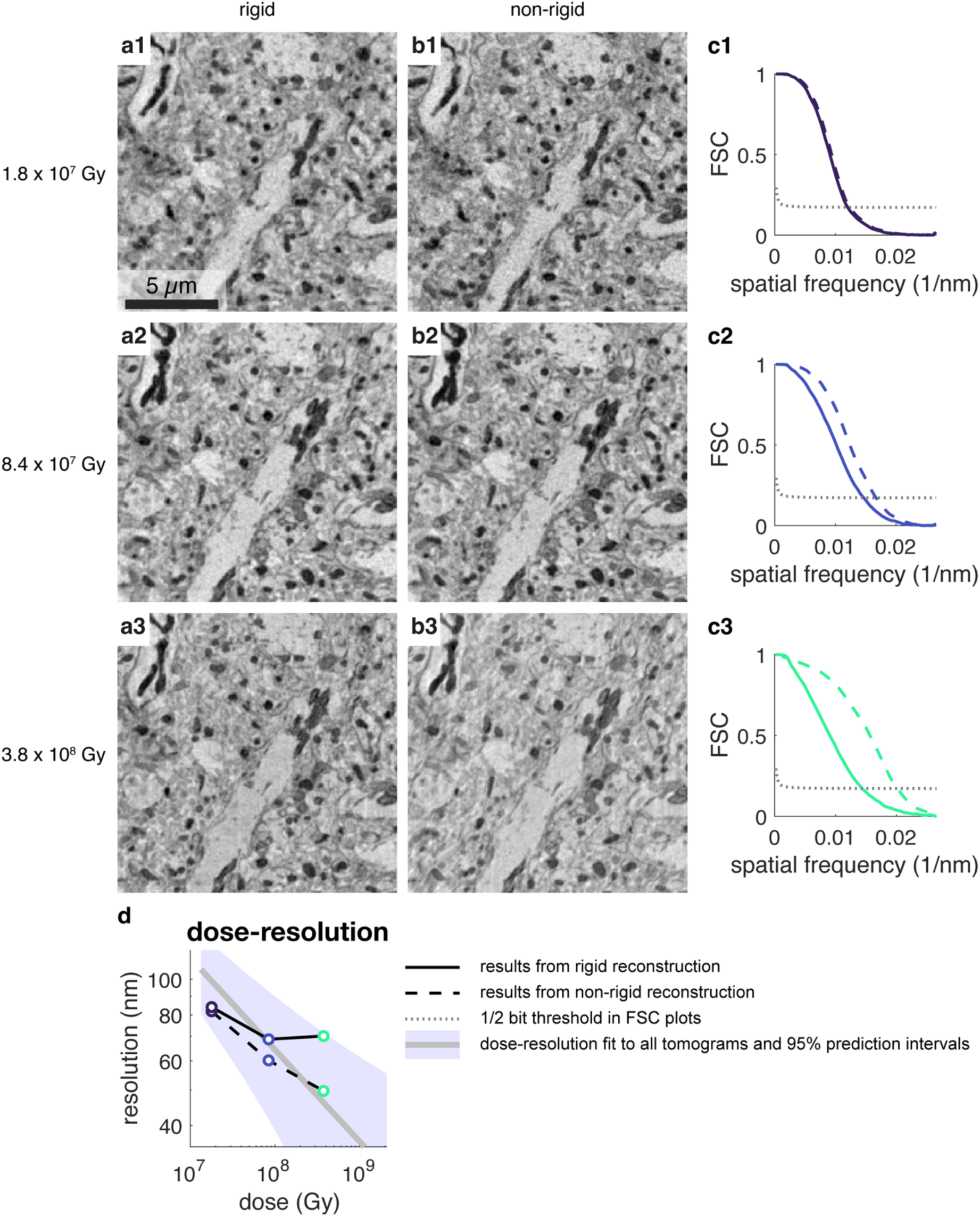
Dose-dependent resolution of the reconstructed tomograms. **(a1-a3)** Example images showing approximately the same region imaged at distinct doses, after a rigid tomographic reconstruction of all subtomograms involved in each case, for a mouse brain EPL sample embedded in resin A (EMbed812). **(b1-b3)** Same images shown in (a1-a3) after non-rigid tomographic reconstruction of the same data. **(c1-c3)** Fourier shell correlation (FSC) for the three tomograms shown in (a,b) and both reconstruction algorithms (solid line: rigid tomographic reconstruction; dashed line: non-rigid tomographic reconstruction). **(d)** Resolution estimated by FSC in (c1-c3) as a function of absorbed dose. Solid and dashed lines indicate rigid and non-rigid tomographic reconstruction, respectively. The solid grey line indicates the fit *FSC_resolution = a/dose^(¼)* for all datasets as described in Fig. 2(d), along the 95% prediction intervals with observation, non-simultaneous bounds (purple shaded area).

**Supp Fig. 2.3:**
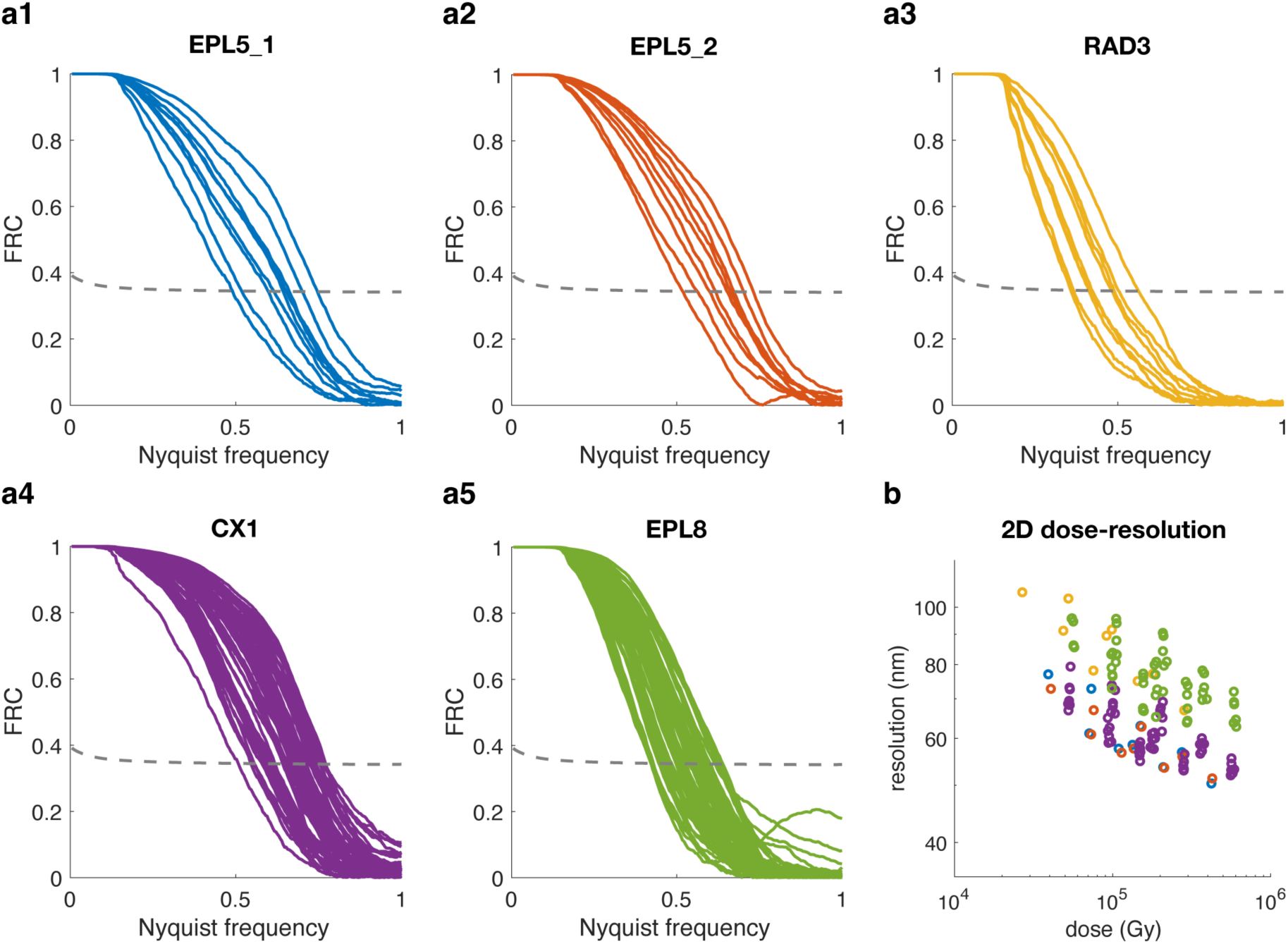
Dose-dependent resolution measurements. **(a1-a5)** Fourier Ring Correlation (FRC) obtained from 2D PXCT images obtained across different X-ray doses on multiple samples from three mouse brain tissue regions (EPL, RAD, CX) embedded in resin A (EMBed812) (EPL5_1, EPL5_2, RAD3, CX1, EPL8) and in resin B (Epon812) (EPL8). **(b)** Resolution estimates for all scans in (a1-a5).

**Supp Fig. 2.4:**
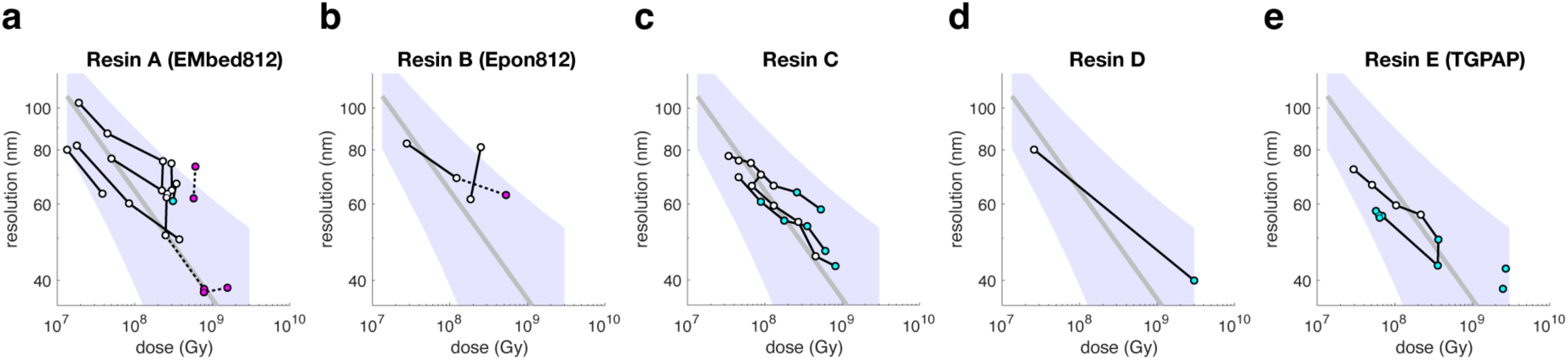
Dose-dependent resolution for samples embedded in all resins. **(a-c)** Fourier shell correlation (FSC) resolution for all tomograms of all samples reported, imaged at different doses, as shown in **Fig. 2d**, split in different subplots by the resin each sample was embedded in. Tomograms (circles) obtained of the same sample at multiple doses form a series and are connected with a black line. The color of the circle indicates whether the tomogram had no radiation-induced damage (white), had contained, globular and compact (<100nm) low-absorbing regions compatible with reconstruction and analysis (cyan) or a diffuse, sparse mass loss (magenta). Examples of the different radiation damage effects are shown in **Fig. 5**. The power function *FSC_resolution = a/dose^(¼)* (grey line) fitted to all white and cyan data points from all resins pooled together along its 95% prediction intervals with observation, non-simultaneous bounds (purple shaded area) is shown in all plots.

**Supp Fig. 4.1:**
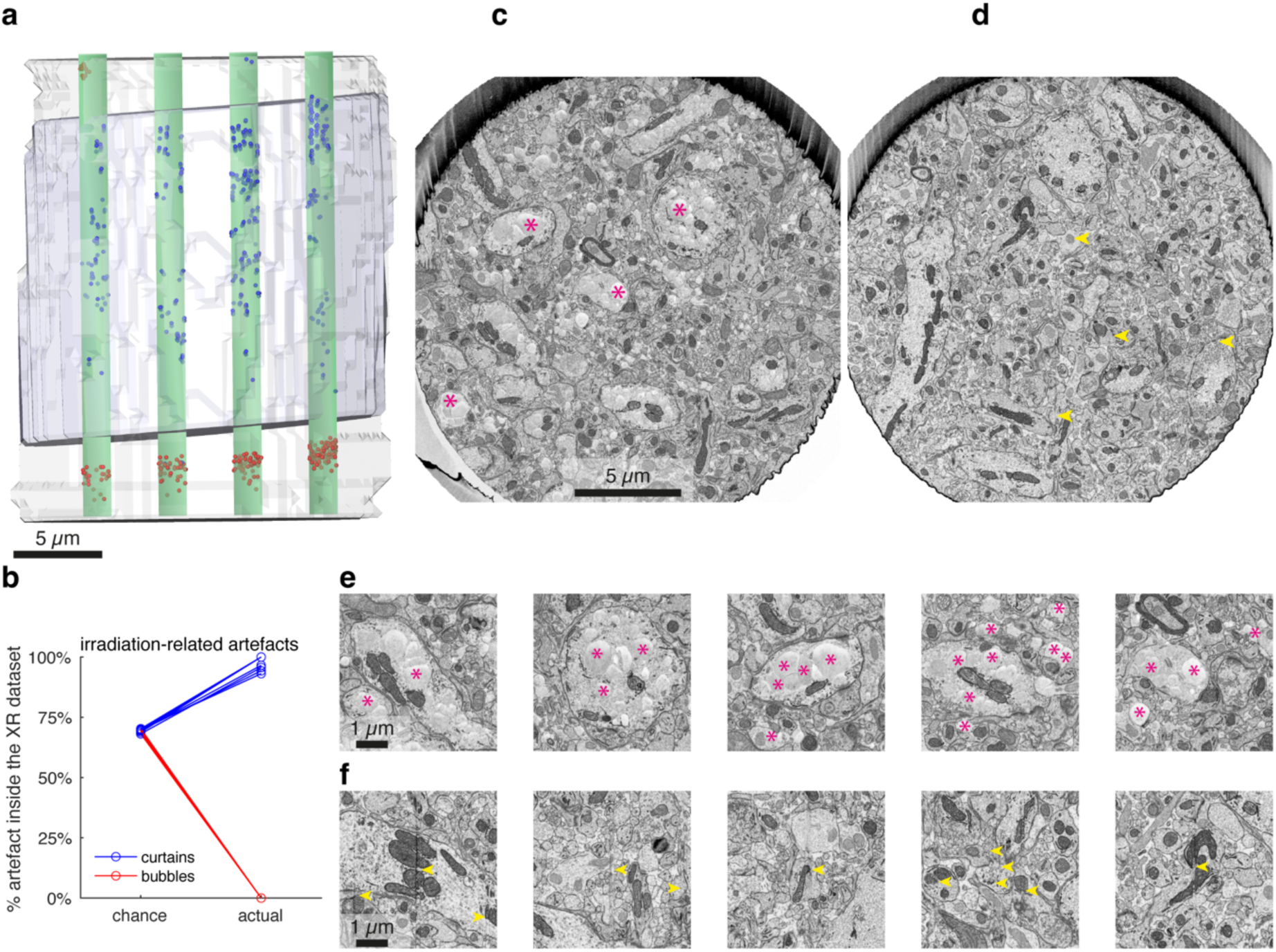
FIB-SEM investigation of ultrastructure. **(a)** 3D render of a mouse brain EPL sample embedded in EMbed812 that was imaged with both PXCT (blue mesh) and FIB-SEM (gray mesh). Note how the FIB-SEM dataset fully contained the region imaged with PXCT. A 4×4 grid of column ROIs (green, only 4 columns visible) was defined and sample defects of two categories (‘curtains’ in FIB-SEM imaging, blue dots and ‘bubbles’, red dots) were annotated throughout the ROIs. For each ROI, its overlap between the two datasets was calculated, and later used to assess the likelihood of artefact distribution by chance vs influenced by X-ray imaging. **(b)** Distribution of the two artefact types inside and outside of the PXCT-imaged region, suggesting that X-ray irradiation triggered both artefacts. Chance level shows, for each ROI, the expected share of artefacts that should have been found in the PXCT-imaged region. Actual level shows the % of curtains (blue) and bubbles (red) found inside the PXCT-imaged region in each ROI. **(c-d)** Cross-section of the FIB-SEM dataset at two heights, showing examples of the artefacts ‘bubble’ (c, magenta asterisks) and ‘curtain’ (d, yellow arrowheads). **(e-f)** Close-up views of the artefacts shown in (c) and (d), respectively.

**Supp Fig. 4.2:**
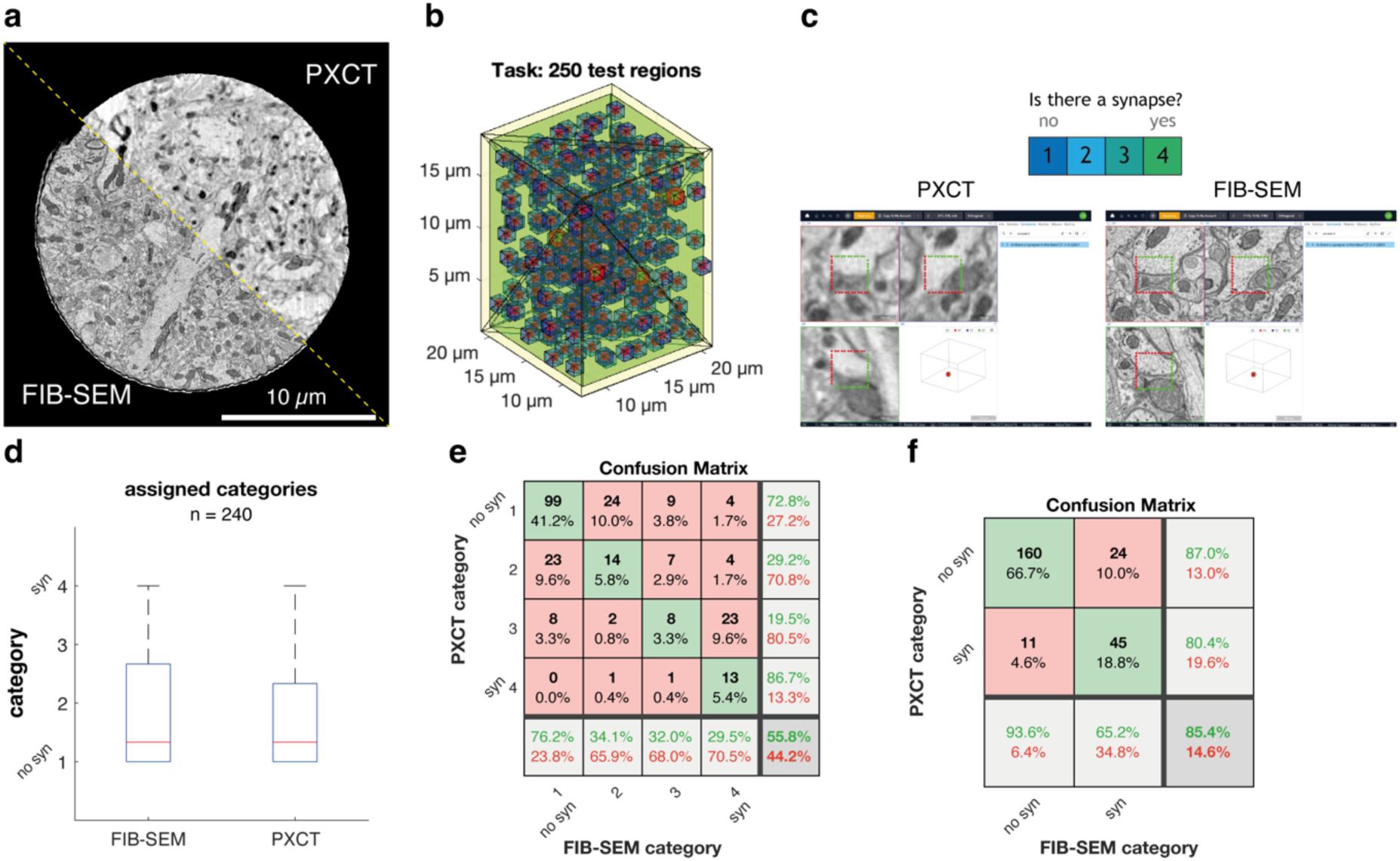
Synapse detection in PXCT. **(a)** Cross-section of a sample imaged by both PXCT and FIB-SEM, showing the same plane viewed by either imaging modality. **(b)** Spatial distribution of 250 non-overlapping regions of interest in the PXCT dataset’s space coordinates, each region occupying 1 µm^3^ (blue) and 10 additional regions used for training (red). **(c)** Synapse ‘captcha’ detection task: a human annotator is presented with each region of interest, displaying either the PXCT or FIB-SEM volume, and must annotate whether the region contains a synapse (score = 4) or it doesn’t (score=1). **(d)** All responses could be obtained from 3 human tracers across 240 regions of interest, and overall the scores distributed equally regardless of imaging modality. **(e-f)** Confusion matrices of synapse detectability in PXCT using FIB-SEM as ground truth, showing all 4 categories (e) and pooling the results into 2 categories (‘synapse’ vs ‘no_synapse’) (f).

**Supp Fig. 5.1:**
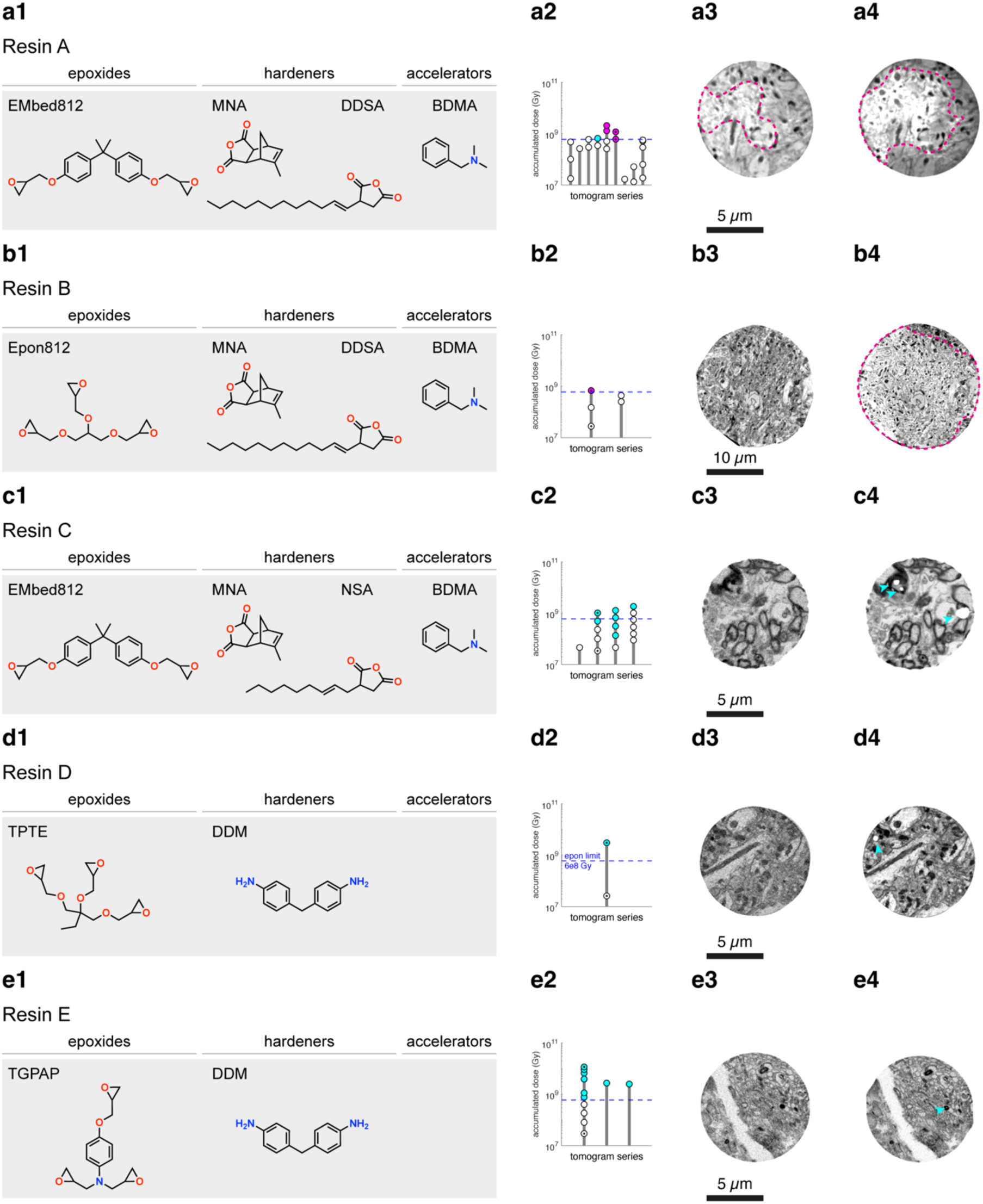
epoxy resins. **(a1)** Monomers employed in the formulation of resin A. **(a2)** Tomogram series acquired with each resin, showing stacked doses of all tomograms for each series, allowing to read in the y-axis the total accumulated dose absorbed by the sample at the end of each tomogram. Colour coding of tomograms is as in Figure 5(d,h), with tomograms displaying localised compact mass loss labelled in cyan, and those displaying widespread mass loss incompatible with high-resolution reconstruction coloured in magenta. **(a3-4)** Representative cross-section of the reconstructed first (a3) and last (a4) tomogram from of a series of tomograms obtained from the same sample location. These tomograms are indicated in (a2) with a black dot. Widespread (magenta) mass loss resulting from extended irradiation is indicated. (b-e) Same as in (a) for the epoxy resins B-E, respectively, used to embed the mouse brain tissue samples reported in this study.

**Supp Fig. 5.2:**
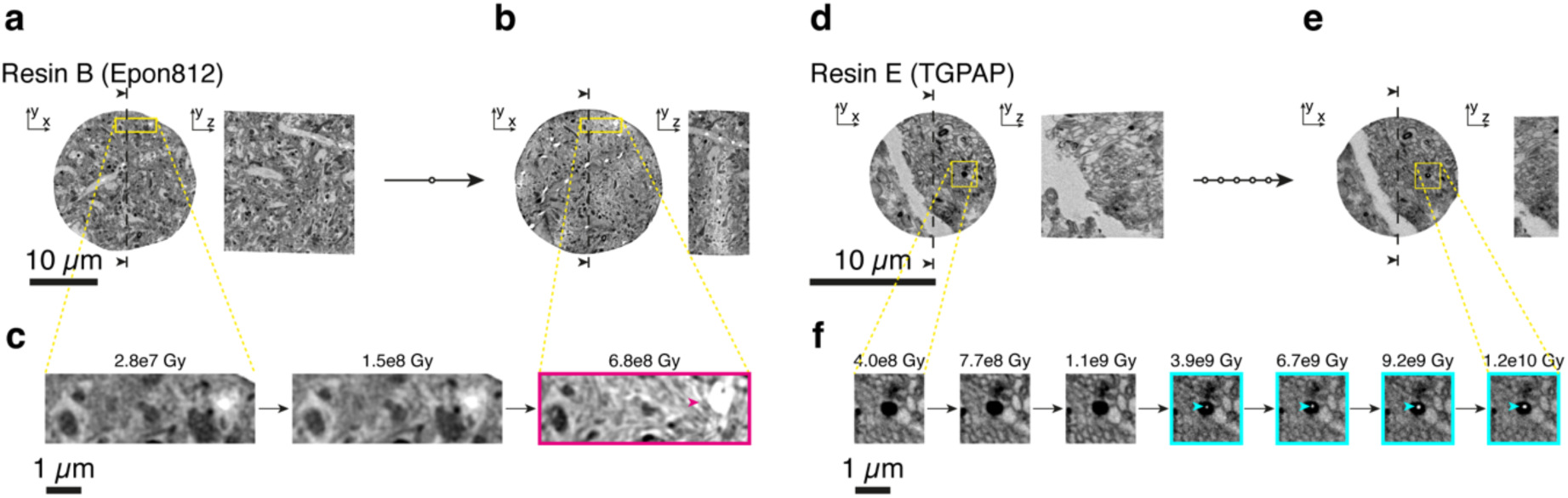
Widespread vs localised compact mass loss effects. **(a-f)** Radiation stress tests performed at cryogenic temperatures under vacuum with a 6.2 keV monochromatic X-ray beam (PSI cSAXS) using ptychographic X-ray tomography on similar samples embedded with resin B (Epon812) (a-c) and resin E (TGPAP) (d-f). The same tissue region was imaged multiple times successively (a-b, d-e) providing a series of tomograms that allowed for both assessing the sample structure and monitoring the radiation dose absorbed. Samples embedded in resin B (Epon812) consistently displayed signs of widespread mass loss (c, magenta arrowheads) after absorbing 6.8e8 Gy.

**Supp Video 5.1.** Radiation-sensitivity of Epon812 and TGPAP embedded stained olfactory bulb tissue.

**Supp Video 5.2.** Tomogram of a TGPAP embedded sample with a resolution of 38 nm.

**Supp Video 5.3.** 3D rendering of tomogram with segmented features.

**Supp Table 1.** Tomogram series.

**Supp Table 2.** Monomers used in all resins.

**Supp Table 3.** Monomer ratios used in each resin.

**Supp Table 4.** Imaging parameters of selected datasets.

**Supp metadata.** Properties of all tomograms. (see Code Availability).

## Materials & Methods

All samples, tomogram series, doses and resolutions of all individual tomograms and hyperlinks to all 3D datasets are listed in **Supp. Table 1**.

### Animals

Animals used in this study were 8-13 week old wildtype mice of C57Bl/6 background of either sex. All animal protocols were approved by the Ethics Committee of the board of the Francis Crick Institute and the United Kingdom Home Office under the Animals (Scientific Procedures) Act 1986.

### Statistics and reproducibility

All images displaying biological features report intuitive examples of features robustly resolved in independent specimens.

### Embedding Resins

In total five different embedding polymers were employed. Each resin originates from the mixture of a set of monomers at specific ratios, and then polymerised at 70°C for ∼72h. Their monomer and putative polymerised structure is reported in **Supp.** Fig. 5**.1**, the monomer identities are listed in **Supp. Table 2**, and the ratios are detailed in **Supp. Table 3**. An overall nomenclature is used -referring to them as ‘A, B, C, D, E’. To enhance readability of the statements, we often simultaneously refer to them with the common name or acronym given to the epoxide monomer (such as ‘Epon812’). Finally, resins ‘A, B, C’ are commonly used in volume electron microscopy and often referred to with the overarching term of ‘hard epon’, which applies to the specific ratios of hardener:resin used ^49^. Accordingly, the term ‘hard epon’ has been used in conjunction with the aforementioned terminology to support readability. In contrast, resin ‘E’ consisting of the epoxy-hardener mixture TGPAP-DDM is referred to as ‘tough’ highlighting its resilience to radiation.

### Tissue preparation

Tissue was prepared as described previously ^11,59^.

#### Dissection

Mice were sacrificed and 600 μm thick sections of brain areas of interest were sliced in ice-cold dissecting buffer (phosphate buffer 65 mM, 0.6 mM CaCl_2_, 150 mM sucrose) with an osmolarity of 300 ± 20 mOsm/L using a LeicaVT1200S vibratome and immediately transferred to ice-cold fixative (either 1% glutaraldehyde or 1.25% glutaraldehyde and 2.5% paraformaldehyde in 150 mM sodium cacodylate buffer pH 7.40, 300 ± 20 mOsm/L). Samples were left in the same fixative overnight, at 4 °C. The fixative was then washed with wash buffer (150 mM sodium cacodylate pH 7.40, 300 ± 20 mOsm/L) three times for 10 min at 4 °C. Overall, samples were kept in an ice-cold, osmolarity-checked buffer.

#### Staining, dehydration and embedding

Slabs were stained with heavy metals using an established ROTO protocol^57^ using an automated tissue processor (Leica EMTP). Briefly, they were first stained with reduced osmium (2% OsO4, 3% potassium ferrocyanide, 2 mM CaCl2 in wash buffer) for 2 h at 20 °C, followed with 1% thiocarbohydrazide (aq) at 60 °C for 50 min, 2% osmium (aq) for 2 h at 20 °C, and 1% uranyl acetate (aq) overnight at 4 °C. On the next day, the samples were further stained with lead aspartate for 2 h at 60 °C. Samples were washed with double-distilled water six times for 10 min at 20 °C between each staining step, except warmer washes before and after TCH (50 °C) and before LA (60 °C).

Samples were then dehydrated with increasing ethanol solutions (75, 90, 2 × 100%), transferred to propylene oxide, and infiltrated with hard Epon mixed with propylene oxide in increasing concentrations (25, 50, 75, 2 × 100%). Finally, samples were transferred into plastic moulds with freshly prepared resin and polymerised individually for 72 h at 70 °C. While final chemical composition is difficult to estimate^75^, overall density of the polymerised resin was 1.24 ± 0.01 g/ml (n = 15 blocks of resin).

#### Hard epon resin protocols

Resins ‘A, B, C’ fall within the broader description of ‘hard epon’ ^49^: an Epon812 or equivalent epoxide with hardeners MNA and a succinic anhydride at a weight ratio of 8:3:5 (**Supp. Tables 2, 3**) ^49^. Their polymerisation is further catalysed with BDMA. All reagents are viscous at room temperature. Resins ‘A’ and ‘B’ were prepared by mixing the epoxide and hardeners at room temperature in a closed container with a magnetic stirrer turning at gentle speed to avoid air being trapped in the viscous liquid. After the solution was homogeneous, the accelerator BDMA was added and stirred similarly until reaching a final homogeneous solution. For Resin ‘C’, each hardener was mixed with the epoxide separately first. Then, the two homogeneous mixtures of epoxy-hardener were stirred similarly as described above, in the presence of the accelerator BDMA (**Supp. Table 3**). *Resin D (TPTE) protocol:* For resin D, 2 g DDM were dissolved at 70°C in 2 ml acetonitrile. Once dissolved, the solution was cooled down and added 2g of the epoxide TPTE (**Supp. Tables 2, 3**) and followed the same polymerisation route as described below for resin E.

#### Resin E (TGPAP, ‘tough’) protocol

After staining and dehydrating the samples as described above, the samples have been immersed in 2 x 100% acetonitrile (EMS-10020-450ML) for 30 min and 60 min, respectively. The TGPAP-DDM resin consists of the tri-functional epoxy resin triglycidyl-p-aminophenol (TGPAP) and the hardener 4,4’-diaminodiphenylmethane (DDM) at weight ratio of DDM : TGPAP = 1:2 (**Supp. Tables 2, 3**)^76^. Because DDM does not mix well with TGPAP at room temperature and in order to facilitate the infiltration, DDM was first dissolved in acetonitrile heated to 70°C and subsequently, TGPAP was added. The samples were incubated in 1:3 resin : acetonitrile for 2 hrs at room temperature, 1:1 resin : acetonitrile for 2-24 hrs at room temperature and subsequently the samples were placed in 1:1 resin : acetonitrile and cured for 12-72 hrs at 80°C. Because the boiling point of acetonitrile is at 82°C, it is important to keep the sample container lid sufficiently open such that the acetonitrile can evaporate during the curing process (**Supp. Table 3**).

#### Sample verification

To verify staining and sample integrity, all samples were imaged with a Zeiss Versa 510 laboratory-based micro-CT^11^. The region of the sample to be targeted was defined based on the microCT imaging results, and the sample was polished and trimmed to expose the layers of interest using a diamond knife (trim 90, Diatome).

#### Pillar preparation

A cylindrical sample was extracted from the Epon-embedded tissue using a 30keV Ga-beam of 13nA on a Zeiss NVision 40 Gallium FIB-SEM at PSI at the targeted locations. The integrated micromanipulator was used to mount the sample on the holder^77^ for the PXCT measurement (Ga-assisted C-depot was used for the fixation on the mount). The pillars were shaped and fine polished with a 30keV Ga-beam of 1.5nA. The TGPAP pillars were first pre-milled with *Preppy* ^78^. Fine polishing was performed with a Ga-FIB-SEM (TFS Helios 600 i, ScopeM) iteratively with currents decreasing from 65 nA to a final 2.5 nA.

### Ptychographic X-ray computed tomography (PXCT)

#### Instrumentation

PXCT was performed using the OMNY instrument (**Fig. 1**) ^43^. The experiment was carried out at the cSAXS beamline of the Swiss Light Source (SLS) at the Paul Scherrer Institute (PSI), Villigen, Switzerland, during beamtimes e18628, e17964, e18604, e18536 and e19533, in 2019-2022 prior to shutdown for the SLS 2.0 multi-bend achromat construction. Details of the components are as follows. Coherent X-rays enter the instrument and pass optical elements that conform to an X-ray lens used to generate a coherent illumination onto the sample of a few µm. These elements are a gold central stop, a Fresnel zone plate (FZP) and an order sorting aperture. The scattered X-rays are measured by a 2D detector (Eiger 1.5M, ^79^). Accurate sample positioning is essential in X-ray ptychography and is achieved by horizontal and vertical interferometers that measure the relative position of the sample with respect to the FZP ^80^. For this, the sample is directly mounted on a reference mirror which is installed on a 3D piezo stage used for scanning the sample in the beam. A rotation stage allows recording projections at different orientations. All measurements were performed at a photon energy of 6.2 keV, corresponding to a wavelength of 2 Å, selected using a fixed-exit double-crystal Si(111) monochromator.

For datasets acquired in all beamtimes except e19533, the FZP used, fabricated by the X-ray nano-optics group at PSI, had a diameter of 220 μm and 60 nm outermost zone width, resulting in a focal distance of 66.0 mm. For datasets acquired during beamtime e19533, we used a FZP from XRNanotech with 250 μm diameter and 30 nm outermost zone width, which had a focal distance of 37.5 mm. Both FZPs featured locally displaced zones, designed to produce an optimal structured illumination for ptychography^81^, and resulted in a beam flux of about 7×10^8^ photons/s. The unscattered beam was blocked by the combination of a 40 μm diameter central stop and a 30 μm diameter order sorting aperture. The FZP was coherently illuminated by using an upstream slit of 20 μm horizontal width, placed at about 12 m downstream of the undulator source, used as a secondary source. The detector was placed at about 7.2 m downstream of the sample.

#### Data acquisition

Samples were placed after the focal spot where the beam had a diameter of approximately 8 μm and 5 μm for the first four beamtimes and for the last beamtime e19533, respectively. In general, ptychographic scans were performed with a field of view that included the full sample size in the horizontal direction, while the scan points were positioned following a Fermat’s spiral trajectory^82^. The dose on each 2D ptychographic projection was adjusted by combining different acquisition settings, namely: (1) the average step size of the scan, ranging from 2.0 to 0.5 μm, and (2) the exposure time at each scan point, ranging from 0.025 to 0.1 s. Reconstructions were performed from areas in the detector ranging between 480 x 480 pixels^2^ to 700 x 700 pixels^2^, resulting in reconstructed pixel sizes of between 41.7 nm and 27.6 nm.

Ptychographic reconstructions of the first four beamtimes were performed with a few hundred iterations of the difference map (DM) algorithm followed by a few hundred iterations of a maximum likelihood (ML) refinement^83^. Ptychographic reconstructions of the last beamtime e19533 were performed with 2 probe modes using 600 iterations of ML.

For the tomography, several projections with equal angular spacing between sample rotations of 0° and 180° were recorded. The number of projections varied between approximately 300 and 1200, providing another approach to adjust the dose imparted on each sample. The phase of the reconstructed projections was used after post-processing alignment and removal of constant and linear phase components^84,85^. A modified filtered back projection was used after aligning the projections using a tomographic consistency approach^84^. The tomograms were computed with a Hann or a RamLak filter for the first four beamtimes and for the last beamtime e19533, respectively. For non-rigid reconstruction, we employed the algorithm described in ^18^.

#### Dose estimation

We estimated the dose as the total energy absorbed by the sample divided by the total mass of the sample. For this the total number of photons absorbed were determined directly from the diffraction patterns, compared with measurements where the beam is not going through the sample. We relied on the linearity of the Eiger 1.5 M detector at maximum measured count rates of 10^5^ photons/(s·pixel) and on its close to 100% efficiency at the used photon energy of 6.2 keV. The sample mass was estimated from the measured volume and the total number of electrons in the sample, which is measured directly by PXCT. For the conversion from the total number of electrons to total mass we made an estimation of the sample composition based on a mixture in equilibrium with all the components used for the embedding resin and the staining, obtaining an average molecular mass estimation of about (1.9±0.1) g/mol, where the error stems from the uncertainty in the sample composition. This estimation is reasonable if we compare it with the values calculated for lipids, 1.80 g/mol^86^, and for a mixture of resin and stain metals with 5 times higher metal content compared to the mixture in equilibrium, 1.99 g/mol. The latter scenario would cause an attenuation through the sample slightly larger than the one measured by X-ray ptychography, providing a reasonable upper limit for the metal content within the sample, and thus for the average molecular mass of the sample material. The error in the determination of the absorbed dose is mostly given by this uncertainty.

The two datasets highlighted in (**Fig. 4**) depict the optimal results obtained when imaging at a dose of 3.8e8 Gy and 2.5e9 Gy, respectively. The parameters employed in the acquisition of these two PXCT datasets in particular are detailed in (**Supp. Table 4**). Detailed description of all reconstructions is provided in (**Supp Metadata**) and a summary can be found in (**Supp. Table 1**).

### FIB-SEM data acquisition

Focused ion beam scanning electron microscopy (FIB-SEM) was carried out using a Crossbeam 540 FIB-SEM with Atlas 5 for 3-dimensional tomography acquisition (Zeiss, Cambridge). The OMNY pin was coated with a 10 nm layer of platinum, mounted horizontally on a standard 12.7 mm SEM stub using carbon cement (LeitC), and coated with a further 10 nm layer of platinum. This method of mounting the pin ensured the cylinder of tissue was positioned in free space and could be reoriented appropriately within the SEM for ion beam milling with consideration to the X-ray dataset. As the sample was cylindrical and tracking marks therefore could not be applied, direct tracking of slice thickness was not possible. Autofocus and autostigmation functions were carried out on an area close to the outer edge of the sample. Electron micrographs were acquired at 8 nm isotropic resolution, using dwell times of 12 µs (C319_EPL1), 6 µs with ×3 line averaging (Y357_3dot), or 8.5 µs (Y357_30um). During acquisition, the SEM was operated at an accelerating voltage of 1.5 kV with 1 nA current (C319_EPL1, Y357_30um) or 500 pA (Y357_3dot). The EsB detector was used with a grid voltage of 1,200 V. Ion beam milling was performed at an accelerating voltage of 30 kV and current of 700 pA. Approximate data acquisition times were: 3 days 7 hours and 3260 slices (C319_EPL1), 2 days 3 hours and 2814 slices (Y357_3dot), 3 days 18 hours and 1936 slices (Y357_30um).

The FIB-SEM dataset was later warped to the ptychography space using Bigwarp^87^. The warped data of the volume also imaged with ptychography was then exported to the ptychography dataset’s space with a voxel size of 9.4 nm in x,y,z in C319_EPL1 and 6.9 nm in Y357_3dot and in Y357_30mu (which is 1/4 of the voxel size of the native ptychography dataset, 37.6 nm and 27.6 nm, respectively). In this way, both datasets could be stored as different layers of a common dataset in webknossos^88^. This setup enabled quick toggling between imaging modalities at any particular location, and simplified the configuration of the synapse detection tasks.

### FIB sample preparation for SLS

The pillars were cut with a 30keV Ga-beam of 13nA on a Zeiss NVision40 FIB-SEM (PSI). They were shaped and fine polished with a 30keV Ga-beam of 1.5nA. The TGPAP pillars were first pre-milled with *Preppy*^78^. Fine polishing was performed with a Ga-FIB-SEM (TFS Helios 600 i, ScopeM) iteratively with currents decreasing from 65 nA to a final 2.5 nA.

### X-ray computed tomography at BM05 at ESRF

The Resin B (Epon812) and Resin E (TGPAP)-embedded samples shown in **Fig. 5** and **Supp Video 5.1** were fixed, stained and embedded as described in the previous sections. For X-ray computed tomography and continuous beam exposure they have been mounted on standard aluminium microtomography holders. Measurements were done in identical conditions for the two samples. The samples were illuminated with a broad bandwidth beam with an average energy of 25 keV, filtered with 0.54 mm of Al. The detector used for imaging was composed of a 23 µm thick LSO scintillator coupled to an infinity-corrected long-working-distance Mitutoyo objective (10x, NA 0.28) and a PCO Edge sCMOS camera. Tomographic scans were recorded with 3000 projections over 360°, using an exposure time of 50 ms per frame. The pixel size was 0.73 µm. Each scan took about three minutes and 45 seconds and between two consecutive scans the sample was continuously exposed to the same beam without imaging for 20 minutes. The Epon embedded sample was exposed for a total of 4 hours, resulting in severe damage as it can be observed in **Fig. 5(b)**. The TGPAP embedded sample was exposed for an additional 1 hour, thus a total of 5 hours, without generating any observable damage. The measurements were performed in air at room temperature. The detector used for imaging was composed of a 23 µm thick LSO scintillator and a PCO Edge sCMOS camera.

### Data analysis

#### FSC analysis

To measure resolution, a custom implementation of Fourier ring or shell correlation analysis was used. For the estimation of the 2D resolution of a projection image, we perform the 2D Fourier ring correlation (FRC) between that image and a second image acquired with identical experimental parameters. We then compare the FRC with a threshold using the 1-bit criterion, which is equivalent to a signal to noise ratio of 0.5 for each of the compared images^51^. This procedure provides an estimation of the half-pitch resolution for each of the individual images. For the estimation of the 3D resolution of a tomographic dataset, we compute the 3D Fourier shell correlation (FSC) of two subtomograms, each computed using half of the tomographic projections. In this way, we obtain two datasets acquired independently, albeit each with double angular sampling. We then compare the FSC with a threshold according to the ½-bit criterion, which corresponds to a signal to noise ratio of 0.4 for the full tomographic dataset.

#### Synapse identification

The synapse identification tasks were performed on the sample for which both FIB-SEM as well as ptychography data were available.

Two synapse identification approaches are presented: a dendrite-centric one and a randomised “captcha”-like detection.

For the dendrite-centric synapse identification task, five dendrites evolving in straight trajectories in distinct directions were chosen and traced (in a skeleton format) from the ground truth FIB-SEM dataset. The dendrites chosen all presented a pale cytoplasm, straight trajectory, no branches and consistent thickness of around 2 µm, to ensure they could be followed in the datasets of both imaging modalities. Since the sample this dataset belongs to was extracted from the external plexiform layer of the olfactory bulb, these dendrites are likely to be lateral (and possibly apical) dendrites of projection neurons (mitral or tufted cells). Each dendrite skeleton was followed three times independently looking at one orthogonal plane only, and every time all features resembling synapses were annotated by seeding single nodes. This operation was performed for both the FIB-SEM as well as the ptychography dataset (n = 725 initial nodes in FIB-SEM, 576 initial nodes in PXCT). All nodes seeded in the FIB-SEM dataset were assumed to be pointing to true synapses. Based on previous studies, synapse density was estimated to be of 1-2 synapse / µm^3 89^. First, we obtained a census of synapses by looking at the FIB-SEM data alone: nodes seeded ≤ 300 nm away from each other were assumed to be pointing to the same synapse, which lead to a total census of 338 synapses in all 5 dendrites. Most synapses (273/338, 81%) were detected on the first pass in the FIB-SEM dataset, and only a small fraction of the final census was only found at the 3rd and last pass (65/338, 19%), consistent with the assumption that synapses are well resolved in the FIB-SEM data. Next, the nodes seeded in the ptychography dataset were matched with the previously defined census of synapses through a similar process: FIB-SEM-validated synapse locations receiving a tag from the ptychography dataset within a 300 nm euclidean distance were marked as ‘detected’. Most detected synapses in X-rays (148/225, 66%) were already detected on the first pass in the ptychography dataset, and only a small fraction were only detected at the 2nd and last passes (77/225, 34%). A number of annotations of putative synapses in the ptychography datasets were not matched to any synapse. These annotations were further used to detect robust confounding factors when annotating synapses. Locations in the dataset receiving ≥ 2 tags within a ≤ 300 nm distance were defined as ‘hot spots’ for confusion. A total of 36 hotspots were detected, and their ultrastructure was revisited in the FIB-SEM dataset. Features providing the confusion were then identified by toggling the view of the ptychography and FIB-SEM dataset in the browser, and ultimately the nature of the feature leading to false positive detection in the ptychography dataset was annotated categorically. After revisiting the ultrastructure of all 36 false positive hotspots, we quantified the prevalence of the different feature categories.

For the “captcha”-like synapse detection task, we generated 250 non-overlapping and randomly located 1*1*1 µm^3^ regions of interest (**Supp Fig. 4.2**). These regions could be displayed as cubes with 3 red faces and 3 green faces in the webknossos environment. The task consisted in determining whether a given region contained a synapse or not, by assigning a confidence score of [1 = clearly not containing a synapse, 2, 3, 4 = clearly containing a synapse]. If the synapse was only partly contained inside the region, it would only be taken into account if it exits the region through a green boundary. Additionally, ten locations known to contain a synapse (extracted from the previous dendrite-centric analysis) were chosen as training data. Finally, the task was encoded within the webknossos ecosystem such as one would be navigated from region to region after each answer was being recorded, and only one dataset (the one being tested each time) would be presented (**Supp Fig. 4.2**). The task, overall, consisted on exploring the X-ray and FIB-SEM appearance of 10 training synapses, then assess the presence of synapses in the 250 regions in the ptychography dataset, and then assess the presence of synapses in the 250 regions in the FIB-SEM dataset. Three independent annotators, all experts in the appearance of synapses when their fine ultrastructure is resolved (e.g. by FIB-SEM) ran the task. The analyses returned therefore a 250 regions x 3 annotators x 2 imaging modalities array of responses, each with a value of either [1, 2, 3, 4]. In some cases, some annotators did not log a response before switching to the next task, which provided a value of 0. Only regions with correct assessments from all 3 annotators were kept, providing a cleaned up array of 240×3×2 responses. By matching region IDs, this array was represented in a 240×6 table and complemented with other regional metadata in additional columns. For every region, the average score from each imaging modality was calculated from all scores given by the three annotators. All 4 possible values were well represented in both imaging modalities (**Supp Fig. 4.2**). At this point, table rows were split into 4 groups according to the average FIB-SEM scores of each region (rounded to the closest integer). This allowed plotting the scores assigned in the ptychography data depending on the score given in the FIB-SEM data at the same region (**Fig. 4(f)**). The average responses of all regions in both modalities were later binned into two categories (average score ≤2 categorised as ‘no synapse’; average score > 2 categorised as ‘synapse’). This allowed extracting a confusion matrix on the detectability of synapses (**Supp Fig. 4.2**).

The segmentation and renderings shown in **Fig. 1c**, **Fig. 5d** and **Supp Video 5.2** have been generated using the image analysis services of https://ariadne.ai.

## Data availability

Source data of the graphs presented in the main figures are provided as a Source Data file. The datasets and major annotations reported in this study are accessible through the associated code repository (see Code Availability).

## Code availability

Analysis code is available from https://github.com/cboschp/ptychoStainedTissue.

A supplementary structured table including metadata supporting the measurements presented is accessible at https://zenodo.org/records/12802475.

Ptychography acquisition and reconstruction code is available from https://www.psi.ch/en/sls/csaxs/software

All datasets can be viewed, annotated and downloaded from the links provided in **Supp Table 1**, also accessible through the aforementioned code repository on github.

## Author contributions

Conceptualization: CB, AD, AAW, ATS

Sample preparation: CB, EM, YZ, AAW

Algorithm development: TA, MG-S, MH

Endstation development: TA, MH, OB, AM, MG-S, GA, AD

PXCT data acquisition: CB, TA, MH, AP, MG-S, AD, AAW, ATS

SXRT data acquisition: AP, PC

FIB-SEM data acquisition: CJP, LC

Data analysis: CB, TA

Manuscript first draft: CB, ATS

Figure preparation: CB, AP, AD, AAW

All authors contributed to the editing of the manuscript

## Acknowledgements

The authors are grateful to the biological research and scientific computing science technology platforms of the Francis Crick Institute. We thank Jessica Waters, Thomas Rees and Jeannine Hess for insights on resin monomers and Ruairi Roberts for expert annotation of synapses. This research was funded in whole, or in part, by the Wellcome Trust (FC001153 and 110174/Z/15/Z to A.T.S.; FC001999 to L.M.C.). For the purpose of Open Access, the author has applied a CC BY public copyright licence to any Author Accepted Manuscript version arising from this submission. We acknowledge the Paul Scherrer Institut, Villigen, Switzerland, and ESRF, Grenoble, France, for provision of synchrotron radiation beamtime at the cSAXS beamline of the Swiss Light Source (proposals 20190654, 20200783 and 20211852) and at the BM05 beamline of the European Synchrotron (proposals IH-LS-3497 and IH-LS-3529), respectively. The authors gratefully acknowledge ScopeM at ETHZ and in particular Joakim Reuteler for their support & assistance in this work. This work was supported by the Francis Crick Institute, which receives its core funding from Cancer Research UK (FC001153 to A.T.S.; FC001999 to L.C.), the UK Medical Research Council (FC001153 to A.T.S., FC001999 to L.C.), and the Wellcome Trust (FC001153 to A.T.S., FC001999 to L.C.). It was also supported by a Physics of Life grant (EP/W024292/1) to A.T.S. and A.P. funded by EPSRC and Wellcome. A.P. acknowledges funding from the European Research Council under the European Union’s Horizon 2020 Research and Innovation Programme (852455). A.A.W. acknowledges funding from the SERI-funded ERCStG *XrayConnectomics*. The work of T.A. is supported by funding from the Swiss National Science Foundation (SNF), Project Number 200021_196898.

